# The polarity protein Yurt associates with the plasma membrane via basic and hydrophobic motifs embedded in its FERM domain

**DOI:** 10.1101/2023.09.28.559937

**Authors:** Clémence L. Gamblin, Charles Alende, François Corriveau, Alexandra Jetté, Frédérique Parent-Prévost, Cornélia Biehler, Nathalie Majeau, Mélanie Laurin, Patrick Laprise

**Affiliations:** Centre de Recherche sur le Cancer, Université Laval, 9 McMahon, Québec, QC, G1R 3S3, Canada.; axe Oncologie du Centre de Recherche du Centre Hospitalier Universitaire de Québec-UL, 9 McMahon, Québec, QC, G1R 3S3, Canada; Département de biologie moléculaire, de biochimie médicale et de pathologie, Faculté de médecine, Université Laval.; Faculty of Biological and Environmental Sciences, University of Helsinki, Helsinki 00790, Finland; Institute of Biotechnology, University of Helsinki, Helsinki 00790, Finland

**Keywords:** Yurt, membrane localization, FERM domain, FERM-adjacent domain, BH motif, epithelial cell polarity

## Abstract

The subcellular distribution of the polarity protein Yrt is subjected to a spatio-temporal regulation in *D. melanogaster* embryonic epithelia. After cellularization, Yrt binds to the lateral membrane of ectodermal cells and maintains this localization throughout embryogenesis. During terminal differentiation of the epidermis, Yrt accumulates to septate junctions and is also recruited to the apical domain. While the mechanisms through which Yrt associates with septate junctions and the apical domain have been deciphered, how Yrt binds to the lateral membrane remains as an outstanding puzzle. Here, we show that the FERM domain of Yrt is necessary and sufficient for membrane localization. Our data also establish that the FERM domain of Yrt directly binds negatively charged phospholipids. Moreover, we demonstrate that positively charged amino acid motifs embedded within the FERM domain mediates Yrt membrane association. Finally, we provide evidence suggesting that Yrt membrane association is functionally important. Overall, our study highlights the molecular basis of how Yrt associates with the lateral membrane during the developmental time window where it is required for segregation of lateral and apical domains.

Summary statement

This study reveals how Yurt associates with membrane lipids in epithelial cells, thereby further illuminating the mechanisms sustaining cell polarization and tissue morphogenesis.

## Introduction

The plasma membrane of epithelial cells is segregated into an apical, a lateral, and a basal domain (Buckley and St Johnston, 2022). This structural organization, which is referred to as epithelial cell polarity, supports many unidirectional functions of epithelia. Epithelial sheets display a selective permeability owing to occluding junctions sealing the intercellular space. In *D. melanogaster*, this function is fulfilled by septate junctions, which are established laterally (Banerjee et al., 2006).

The protein Yurt (Yrt) plays a crucial role in segregating apical and lateral domains in *D. melanogaster* embryonic epithelia (Laprise et al., 2006; Laprise et al., 2009). In addition, Yrt controls septate junction permeability (Laprise et al., 2009). Yrt encloses a Four-point-one, Ezrin, Radixin, Moesin (FERM) domain at its N-terminus (Hoover and Bryant, 2002; Tepass, 2009). Yrt also contains a FERM-adjacent (FA) domain that is present in a subset of FERM family members (Baines, 2006; Tepass, 2009). The FERM and FA domain of Yrt both contribute to its oligomerization and function (Gamblin et al., 2014; Gamblin et al., 2018). The FA domain of Yrt is followed by a region lacking known protein domains designated as the variable region (VR), which ends with a putative PSD-95, DLG1, ZO-1 (PDZ) domain binding motif (PBM) (Laprise et al., 2006).

After cellularization, Yrt primarily localizes to the lateral membrane (Laprise et al., 2006). During terminal differentiation of epithelial tissues, Yrt accumulates at forming septate junctions and is recruited apically (Laprise et al., 2006; Laprise et al., 2009). While the adhesion protein Neuroglian (Nrg) recruits Yrt at septate junctions (Bieber et al., 1989; Laprise et al., 2009), the apical localization of Yrt depends on its direct interaction with Crumbs (Crb) (Laprise et al., 2006). We thus gained insights into the molecular basis of Yrt association with septate junctions and the apical domain, but it remains to be determined how Yrt associates with the lateral membrane basal to septate junctions or prior to their formation. Furthermore, the functional importance of Yrt outside of septate junctions or the apical Crb complex remains unexplored. In the present study, we performed an *in vivo* structure-function analysis to define the molecular basis of Yrt association with the lateral membrane in the embryonic ectoderm prior to septate junction genesis and assessed the functional importance of Yrt lateral localization.

## Results and discussion

### The FERM domain targets Yrt laterally *in vivo*

The FERM domain protein Yrt primarily associates with the lateral domain prior to septate junction formation in the embryonic ectoderm of stage 10-11 *D. melanogaster* embryos [(Laprise et al., 2006); Fig. 1A, B]. Similarly, immunofluorescence performed with anti-FLAG antibodies showed a lateral enrichment for FLAG-tagged full length (FL) Yrt at these developmental stages ((Gamblin et al., 2014); Fig. 1A, C, M), whereas FLAG-green fluorescent protein (GFP) was distributed throughout cells (Fig. 1D; anti-Lethal giant larvae [Lgl] antibodies were used to visualize the lateral membrane to quantify the membrane-to-cytoplasmic ratio of the GFP signal and other cytoplasmic mutant proteins). This indicates that FLAG-Yrt represents a relevant model to perform a structure-function analysis exploring the molecular basis of Yrt localization. We first investigated the requirement of the FERM domain, the FA domain, and the VR (Hoover and Bryant, 2002; Laprise et al., 2006; Tepass, 2009) for Yrt membrane localization. We found that truncated Yrt proteins made of the FERM domain alone or a combination of the FERM and FA domains showed a similar distribution to FLAG-Yrt FL (Fig. 1A, E, F, M), whereas removal of the FERM domain totally abrogated association of Yrt with the lateral membrane (Fig. 1A, G, M). In contrast, deletion of the FA domain had no significant impact on Yrt lateral distribution (Fig. 1A, H, M). Of note, FLAG-Yrt^ΔFA^ and FLAG-FERM decorated the apical free surface in addition to the lateral membrane (Fig. 1E, H). This is consistent with our previous findings showing that phosphorylation of the FA domain by atypical protein kinase C (aPKC) is required for apical exclusion of Yrt (Gamblin et al., 2014). Moreover, the phosphomimetic FLAG-Yrt^5D^, in which the aPKC phosphorylation sites in the FA domain were mutated to aspartate (D) residues, showed reduced membrane-to-cytoplasmic ratio (Fig. 1A, I, M; (Gamblin et al., 2014)). Finally, truncations made of the FA domain combined with the VR, the VR alone, or the VR amputated from the putative PBM did not show clear enrichment at the lateral domain of cells (Fig. 1A, J-M). Together, these data demonstrate that the FERM domain is necessary and sufficient for Yrt membrane targeting and suggest that the FA domain and the VR have a limited, if any, role in promoting Yrt lateral localization. However, the FA domain likely acts as a phospho-dependent negative regulatory unit controlling Yrt association with the plasma membrane.

**Fig 1.**
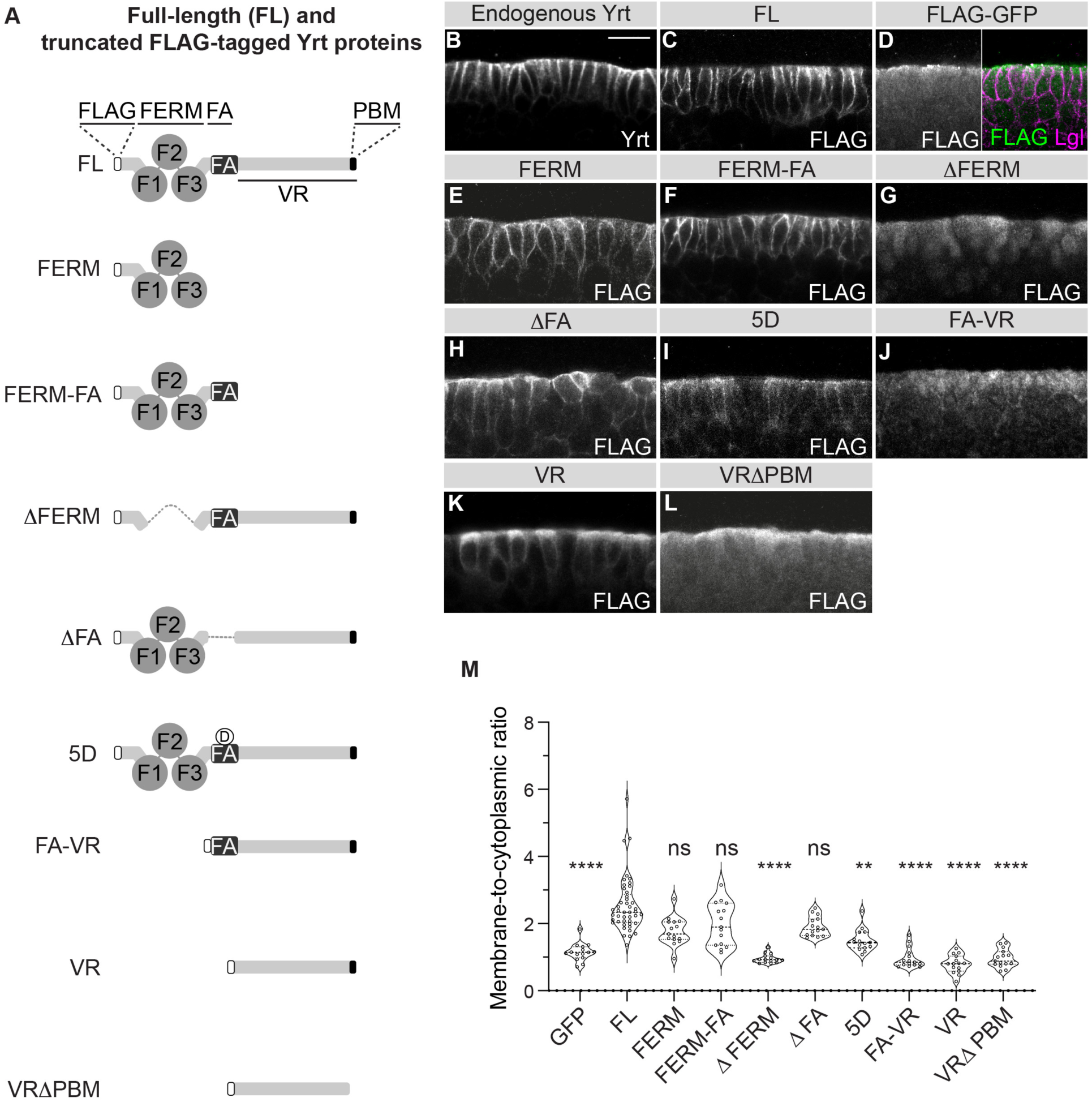
The FERM domain of Yrt is necessary and sufficient for cortical targeting. **(A)** Schematic representation of the FLAG-tagged proteins used in this study. **(B)** Stage 11 wild-type embryo stained for Yrt. The picture depicts a portion of the ventral ectoderm. Scale bar represents 10 µm. **(C-L)** FLAG and Lgl co-immunostaining in stage 10-11 embryos expressing indicated FLAG-tagged constructs. Lgl staining was used to localize the membrane when needed (only shown in D). **(M)** Violin plot showing the quantification of the membrane-to-cytoplasmic ratio of each construct expressed in embryonic ectoderm (bold dotted line = median ; dotted lines = quartiles). Data represents 15 non-adjacent cells per condition (5 cells per embryo from 3 different biological replicates). Kruskal-Wallis test followed by Dunn’s multiple comparison test compare different groups to FLAG-Yrt FL. Non-significant (ns) > 0.05 ; ** *P* = 0.0024 ; **** *P ≤* 0.0001.

### Yrt binds to negatively charged phospholipids and putatively directly associates with the plasma membrane

The fact that FLAG-Yrt^5D^ displays impaired membrane association led us to suggest that the negative charges of the phosphomimetic residues generate electrostatic repulsion with negatively charged membrane phospholipids, thereby interfering with a putative direct association of Yrt FERM domain with the plasma membrane. In support of this hypothesis, the FERM domain of several proteins can directly associate with phospholipids such as phosphatidylinositol 4,5-bisphosphate [PI(4,5)P2] via electrostatic interactions (Ehret et al., 2023). Similarly, purified Yrt FERM domain fused to GST showed affinity for phosphatidylinositol phosphates immobilized on a membrane without clear discrimination between PI(3)P, PI(4)P, PI(5)P, PI(3,4)P2, PI(3,5)P2, PI(4,5)P2, or PI(3,4,5)P3 (Fig. 2A). The negatively charged phosphate group(s) on phosphatidylinositol phosphates is/are required for binding, as GST-FERM only faintly associated with unphosphorylated Phosphatidylinositol (PI) (Fig. 2A). GST-FERM also associated with Phosphatidic Acid (PA) and Phosphatidylserine (PS) (Fig. 2A). However, the FERM domain of Yrt did not interact with Lysophosphatidic Acid (LPA), Lysophosphocholine (LPC), Phosphatidylethanolamine (PE), Phosphatidylcholine (PC), or Sphingosine-1-phosphate (S1P) (Fig. 2A). To test the specificity of our assay, we used GST alone or GST fused to the VR portion of Yrt, which does not associate with the membrane (Fig. 1K), as negative controls. Strikingly, these proteins did not show evident lipid binding (Fig. 2A). We also used the PI(4,5)P2 probe GST-Grip that mainly associated with PI(4,5)P2 (Fig. 2A), as expected (Barnett et al., 2019), thereby showing the validity of our assay. Together, these results show that the FERM domain of Yrt can directly associate with selected lipids such as phosphatidylinositol phosphates but is not a promiscuous lipid-binding protein. This indicates that Yrt may bind the plasma membrane by interacting with charged phospholipids. To test this premise, we decreased the level of plasma membrane PI(4,5)P2 in S2 cells using ionomycin, which leads to PI(4,5)P2 hydrolysis via phospholipase C activation (Sparks et al., 2011). In addition, we used the PI3-kinase inhibitor wortmannin and the PI4-kinase inhibitor phenylarsine oxide (PAO) to decrease the level of some phosphatidylinositol phosphate species phosphorylated at position 3 and/or 4 (Dong et al., 2015; Wymann et al., 2003). FLAG-Yrt was confined at the periphery in cells treated with the vehicle DMSO, whereas it also localized in the cytoplasm in presence of ionomycin or wortmannin and PAO (Fig. 2B-I). Residual membrane association in presence of these inhibitors is consistent with the fact that Yrt can interact with most phosphatidylinositol phosphates and other lipids (Fig. 2A). This also likely explains why PAO and wortmannin have a weak impact on Yrt localization when used separately (Fig. 2E, F, I). Together, these data suggest that Yrt directly associates with phosphatidylinositol phosphates at the lateral membrane in epithelial cells prior to septate junction assembly in *D. melanogaster* embryos, a developmental time window where Yrt contributes to maintaining apical-basal polarity (Laprise et al., 2009).

**Fig 2.**
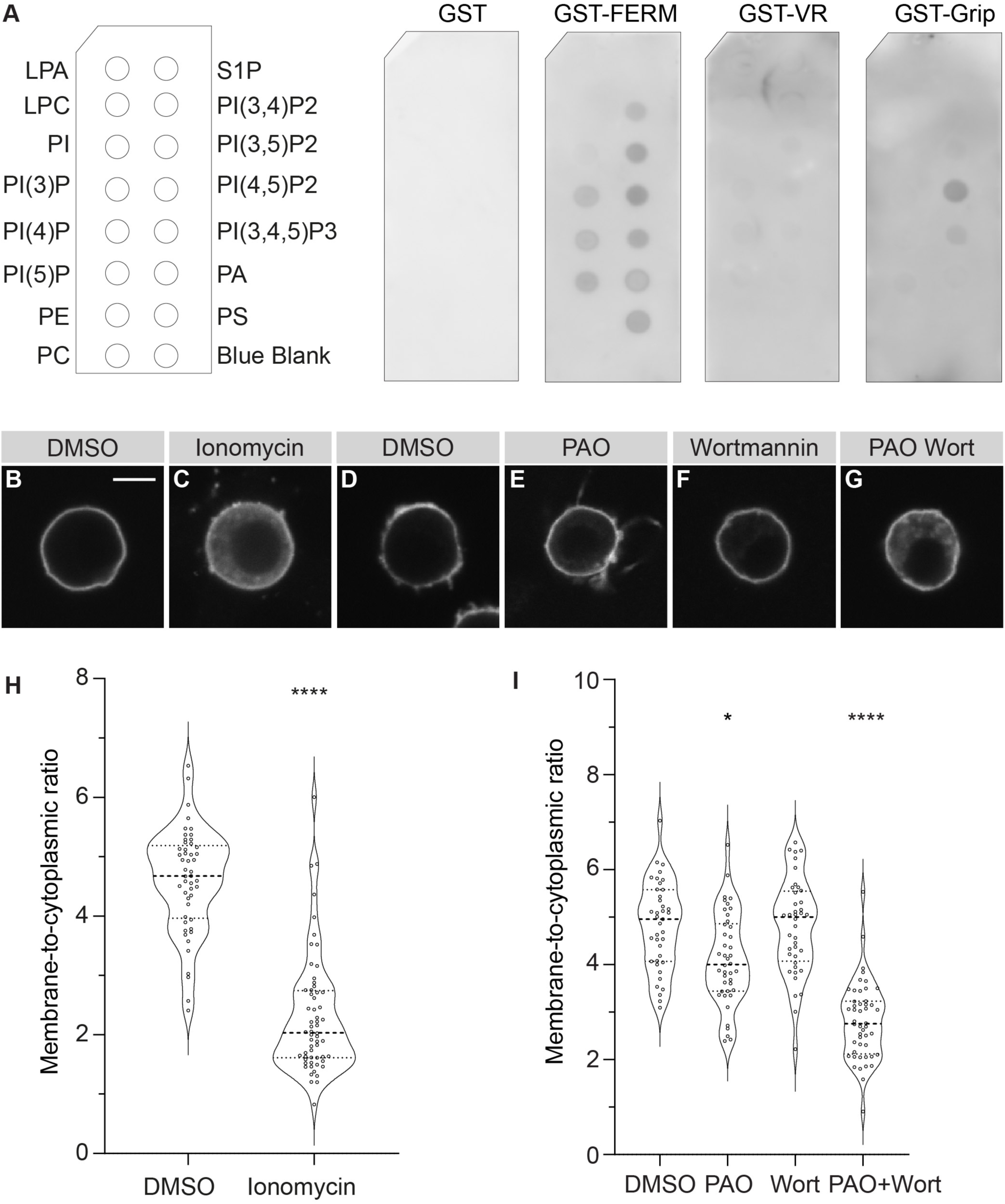
The FERM domain of Yrt associates with negatively charged phospholipids. (**A**) PIP strip membranes were incubated with GST or indicated GST fusion proteins. (**B-F**) S2 cells were transfected with FLAG-Yrt and treated with indicated compounds (E-G). Cells were then stained for FLAG. Scale bar represents 5 μm. (**H-I**) Violin plots showing the quantification of the membrane-to-cytoplasmic ratio in FLAG-Yrt expressing S2 cells presented in B-G (bold dotted line = median ; dotted lines = quartiles). (**H**) Mann-Whitney test, two-tailed with statistical significance set at *P* < 0.05, compares the difference between cells treated with DMSO (n= 49 cells from 3 different experiments) or ionomycin (n=60 cells from 3 different experiments). (**I**) Kruskal-Wallis test followed by Dunn’s multiple comparison test compares each treated group (PAO : n=42 cells ; wort : n=40 cells ; PAO+wort : n=48 cells ; all from 3 different experiments) to vehicle (DMSO ; n=39 cells from 3 different experiments). Non-significant (ns) > 0.05 ; * *P*=0.0265 ; **** *P* ≤ 0.0001.

### Positively charged amino acids within the FERM domain target Yrt to the membrane

Positively charged basic and hydrophobic (BH) amino acid motifs bind membrane phospholipids via electrostatic interactions and contribute to membrane association of selected polarity proteins (Bailey and Prehoda, 2015; Brzeska et al., 2010). Analysis of Yrt sequence revealed the presence of two BH motifs within the FERM domain (regions of the black curve extending above the threshold of 0.6 in Fig. 3A (Bailey and Prehoda, 2015)), which we referred to as BH1 and BH2. Specifically, the BH1 and BH2 motifs span from amino acids 125-130 and 283-293, respectively. *In silico* analysis showed that replacement of the positively charged lysine (K)124, arginine (R)132, and K134 by negatively charged D residues abolished the BH properties of the BH1 motif (Fig. 3A, orange curve). Strikingly, FLAG-Yrt^K124D,R132D,K134D^ (FLAG-Yrt^BH1K/R>D^ hereafter) displayed a strongly decreased membrane-to-cytoplasmic ratio (Fig. 3B, D). Similarly, replacement of K285 and K288 by D residues within the BH2 motif (FLAG-Yrt^BH2K>D^) decreased Yrt membrane enrichment, but the effect on Yrt localization resulting from BH2 mutations was less pronounced than BH1 mutations (Fig. 3C, D). Of note, the BH2K>D mutant has a BH score close to the threshold of 0.6 (Fig. 3A, purple curve). This potentially explains the weak impact of the BH2K>D mutations on Yrt membrane localization. To test this hypothesis, we additionally substituted the last two K residues of the BH2 motif by alanine (A) residues (K292 and K293; FLAG-Yrt^BH2K>D,K>A^). These mutations strongly decreased the BH score (Fig. 3A, blue curve) and led to a strong reduction of Yrt membrane-to-cytoplasmic ratio in S2 cells, which was more significant than the delocalization of FLAG-Yrt^BH2K>D^ and similar to FLAG-Yrt^BH1K/R>D^ (Fig. 3E-J). Together, these results demonstrate that BH1 and BH2 both contribute to target Yrt at the membrane and likely do so via electrostatic interactions. Interestingly, these motifs are conserved in vertebrate orthologs of Yrt (*i.e.* EPB41L4B and EPB41L5 (Tepass, 2009); Fig. S1), thereby suggesting that direct membrane binding is an evolutionarily conserved property of these proteins.

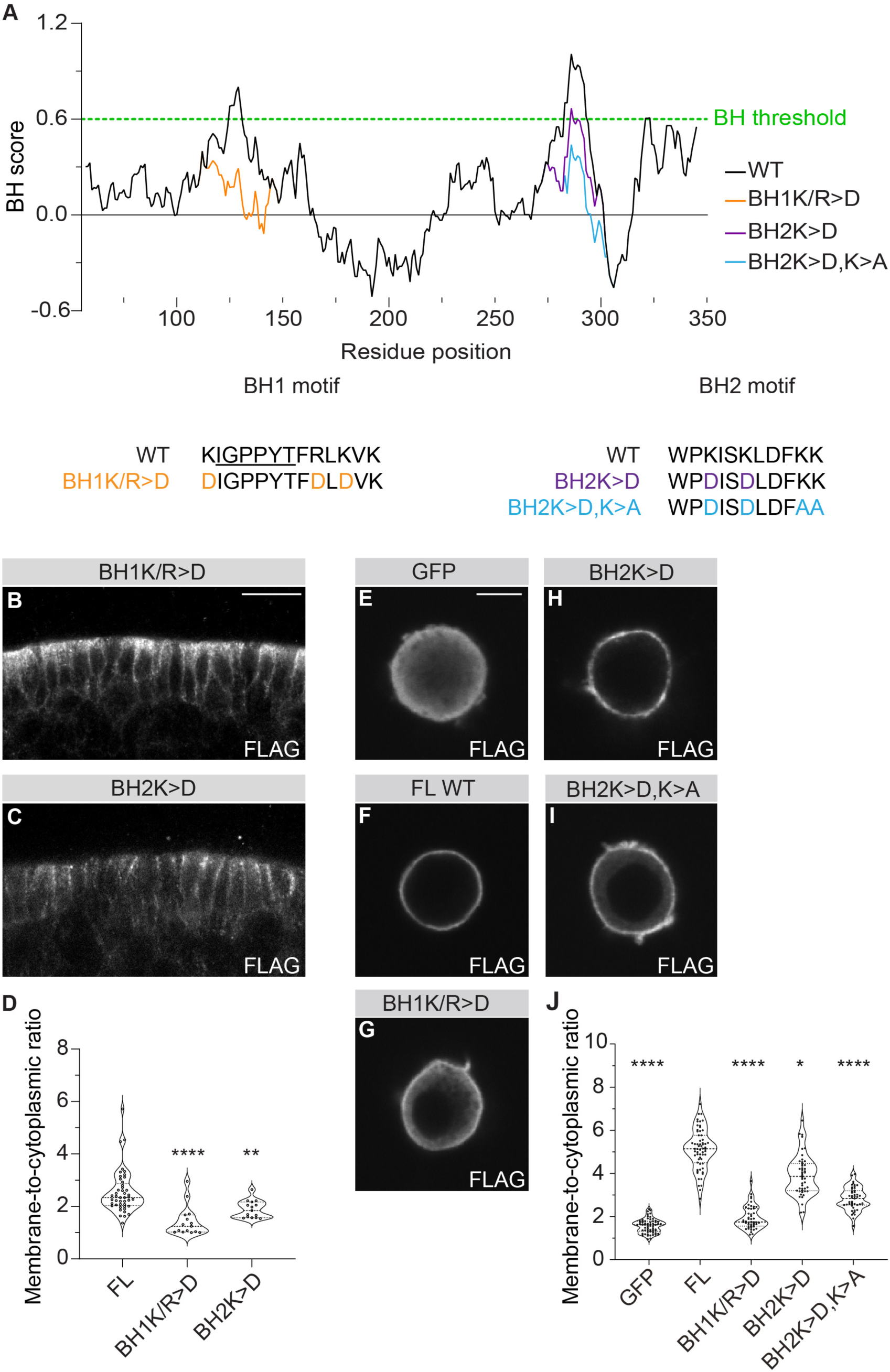
(**A**) The black line in the upper panel shows the BH score (Y axis) along the wild-type sequence of the Yrt FERM domain (X axis). Stretches of amino acids above the threshold of 0.6 are considered BH motifs, which are underlined in the lower panel. The latter also displays mutations introduced in the BH1 (orange) and the BH2 (purple and blue) motifs. The BH score of the mutant sequences is also shown in the upper panel according to the same color code. (**B-C**) Immunostainings performed with anti-FLAG antibodies in stage 10-11 embryos expressing indicated FLAG-tagged constructs. Scale bar represents 10 μm. (**D**) Violin plot showing the quantification of the membrane-to-cytoplasmic ratio for FLAG-Yrt FL, FLAG-Yrt^BH1K/R>D^ and FLAG-Yrt^BH2K>D^ expressed the embryonic ectoderm (bold dotted line = median ; dotted lines = quartiles). Data represent 15 non-adjacent cells per condition (5 cells per embryo from 3 different biological replicates). (**E-G**) Immunostainings performed with anti-FLAG antibodies in S2 cells expressing indicated FLAG-tagged constructs. Scale bar represents 5 μm. (**J**) Violin plot showing the quantification of the membrane-to-cytoplasmic ratio for FLAG-GFP (n=62 cells from 4 different experiments), FLAG-Yrt FL (n=62 cells from 4 different experiments), FLAG-Yrt^BH1K/R>D^ (n=47 cells from 3 different experiments), FLAG-Yrt^BH2K>D^ (n=46 cells from 3 different experiments) and FLAG-Yrt ^BH2K>D,K>A^ (n=47 cells from 3 different experiments) expressed in S2 cells (bold dotted line = median ; dotted lines = quartiles). (**D, J**) Kruskal-Wallis test followed by Dunn’s multiple comparison test compare each group to FLAG-Yrt FL. * *P* = 0.0200; ** *P* = 0.0042; **** *P* ≤ 0.0001.

### Yrt lateral localization is functionally important

Yrt overexpression is associated with a gain-of-function phenotype resulting from altered aPKC and Crb functions (Biehler et al., 2021; Gamblin et al., 2014). Consequently, FLAG-Yrt expression causes epithelial morphogenesis defects and a fully penetrant embryonic lethality phenotype (Gamblin et al., 2014; Gamblin et al., 2018). Similarly, the membrane-associated protein FLAG-FERM-FA is fully lethal (Fig. 4A). FLAG-Yrt^BH1K/R>D^ and FLAG-Yrt^BH2K>D^, which show cytoplasmic delocalization with residual membrane association (Fig. 3B-D), showed a reduced ability to cause embryonic lethality (Fig. 4A). Strikingly, the cytoplasmic truncations FLAG-Yrt^ΔFERM^, FA-VR, VR, and VRΔPBM did not cause any lethality (Fig. 1G, J-M; 4A), although these constructs showed a similar expression level to construct causing lethality (Fig. 4B). These results suggest that the gain-of-function phenotype caused by Yrt overexpression reflects its lateral levels, indicating that Yrt plays an important function at this subcellular location. Alternatively, our data may reflect the functional importance of the FERM domain, as all mutant or truncated proteins that are phenotypically silent or that cause a weak lethality phenotype enclose an impaired FERM domain. To distinguish between these two possibilities, we used FLAG-Yrt^5D^, which contains phosphomimetic mutations on aPKC target sites within the FA domain while comprising an intact FERM domain (Gamblin et al., 2014). FLAG-Yrt^5D^ displayed reduced membrane levels concomitant with cytoplasmic accumulation and is expressed at a similar level to wild type FLAG-Yrt (Fig. 1I, 4B; (Gamblin et al., 2014)). This is in line with our previous observation that ectopic localization of aPKC to the lateral domain displaces Yrt from the membrane (Biehler et al., 2021). Our data thus further illuminate how Yrt function is antagonized by aPKC, which likely represses Yrt association with membrane phospholipids via electrostatic repulsion (this study) while also preventing Yrt oligomerization and binding to Crb (Gamblin et al., 2018). Importantly, expression of FLAG-Yrt^5D^ had a modest impact on embryo survival with an 80% hatching rate (Fig. 4A). This is consistent with the idea that Yrt needs to associate with the lateral membrane to be fully functional. However, this result must be interpreted with caution since FLAG-Yrt^5D^ displays impaired oligomerization, which also contributes to Yrt function (Gamblin et al., 2018). To identify a Yrt mutant protein with impaired membrane association but showing intact oligomerization, we tested whether mutation of the BH1 or BH2 motifs impacts Yrt oligomerization. We found that FLAG-tagged Yrt^BH2K>D^ and Yrt^BH2K>D,K>A^ co-precipitated with HA-Yrt similar to wild-type FLAG-Yrt (Fig. 4C). This shows that these proteins maintained their ability to oligomerize. In contrast, FLAG-Yrt^F281A, W283A^ failed to associate with HA-Yrt (Fig. 4C), as predicted by our previous work (Gamblin et al., 2018). FLAG-Yrt^BH1K/R>D^ also showed impaired association with HA-Yrt (Fig. 4C). We also expressed a construct made of GFP fused to the transmembrane domain and cytoplasmic tail of Crb with FLAG-Yrt^BH1KR>D^, FLAG-Yrt^BH2K>D^ or FLAG-Yrt^BH2K>D,K>A^ to test whether mutation of the BH motifs modifies the association with Crb, which supports some Yrt functions such as apical constriction and maintenance of the apical/lateral membrane ratio (Biehler et al., 2021; Laprise et al., 2006; Laprise et al., 2010). FLAG-Yrt proteins with an impaired BH2 motif associated with GFP-Crb like their wild-type counterpart (Fig. 4D). In contrast, the FLAG-Yrt^L236R^ showed weak binding to Crb, as expected (Gamblin et al., 2018; Ohata et al., 2011), so did FLAG-Yrt^BH1KR>D^ (Fig. 4D). Given that overexpression of FLAG-Yrt^BH2K>D^, which show reduced membrane localization while maintaining its ability to oligomerize and interact with Crb, is associated with reduced lethality as compared to overexpression of wild-type FLAG-Yrt or membrane-bound FLAG-FERM-FA (Fig. 4A), we conclude that association of Yrt with the lateral membrane is most likely important to sustain certain of its functions.

**Fig 4.**
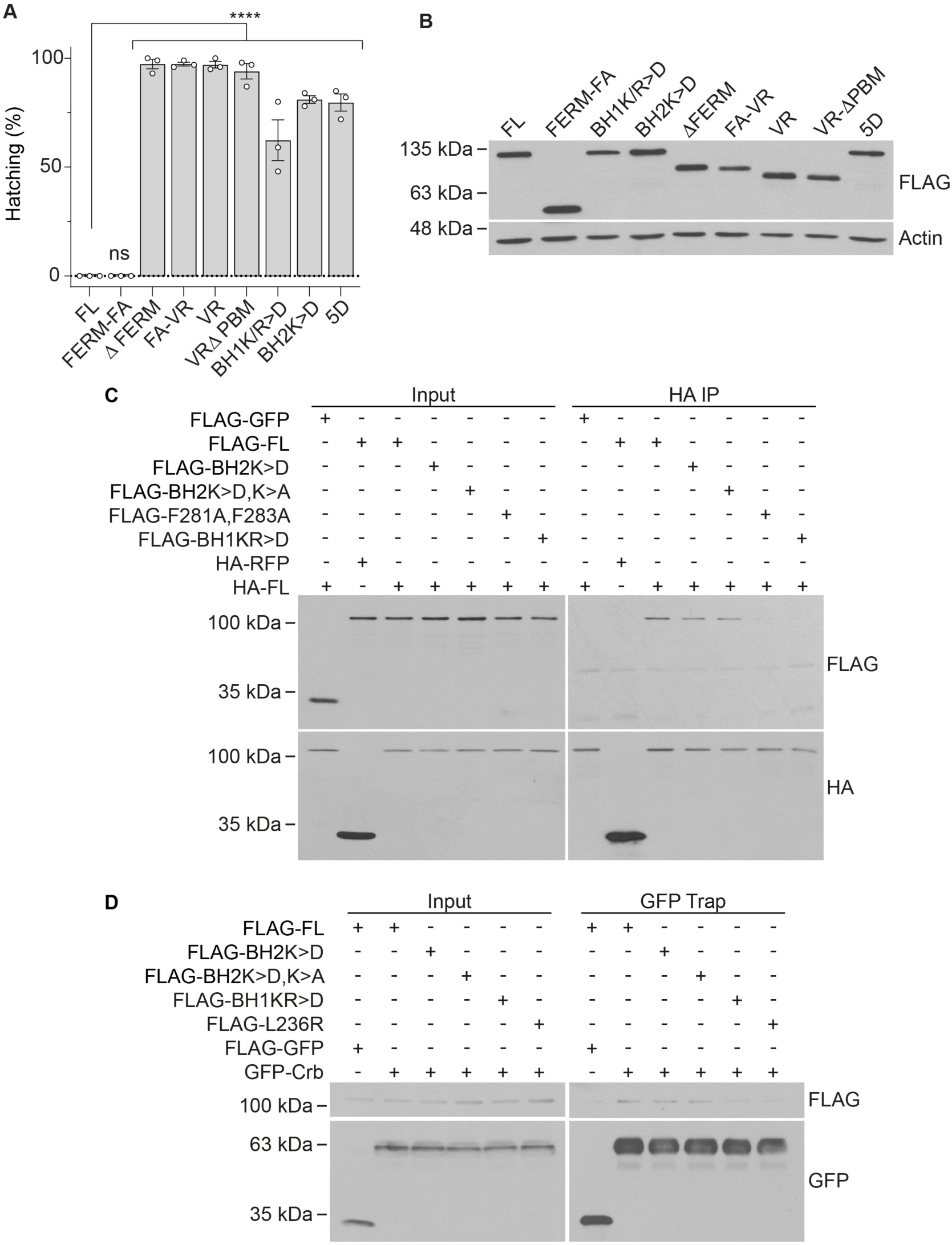
Yrt association with the lateral membrane is functionally important. (**A**) Histogram showing the hatching rate, calculated in percentage, of embryos expressing indicated FLAG-tagged Yrt proteins. Data are shown as means ± SEM, and statistical significance was assessed by using a two-tailed Fisher’s exact test (FL: *n* = 269; FERM-FA: *n* = 234; ΔFERM: *n* = 270; FA-VR: *n* = 233; VR: *n* = 269; VRΔPBM: *n* = 235; BH1K/R>D: *n* = 237; BH2K>D: *n* = 265; 5D: *n* = 247 ; all from 3 different experiments). The significance threshold was adjusted in regards of the multiple comparisons with a Bonferroni correction (8 comparisons). Non-significant (ns) > 0.00625 ; **** P ≤ 0.0001. (**B**) Western blot showing the relative expression of indicated FLAG-tagged constructs in stage 11-12 embryos. Actin was used as loading control. (**C**) S2 cells were transfected with indicated FLAG- and HA-tagged constructs. Immunoprecipitations with anti-HA antibodies were then performed. (**D**) S2 cells expressing GFP-tagged transmembrane and cytoplasmic domain of Crb and indicated FLAG-tagged Yrt proteins were homogenized and submitted to GFP-Trap immunoprecipitations.

Overall, our study shows that BH motifs within the FERM domain contribute to the localization of Yrt at the lateral membrane putatively through direct association with negatively charged plasma membrane phospholipids, including phosphatidylinositol phosphates. Mutations of the positively charged residues contained in BH motifs attenuate the Yrt gain of function phenotype, thereby arguing that Yrt association with the lateral membrane is functionally important. Yrt could thus be an important interactor of phosphatidylinositol phosphates allowing these molecules to perform their known role in establishing and maintaining apical-basal polarity (Gassama-Diagne et al., 2006; Krahn, 2020; Martin-Belmonte et al., 2007).

## Materials and methods

### Molecular biology, transgenic fly lines, and *in vivo* expression of transgenes

The In-Fusion cloning kit (Takara Bio) was used according to manufacturer’s guideline to clone cDNAs into pUASTattB (kindly provided by K. Basler, University of Zurich) and pGEX-6P-2 (GE Healthcare Life Sciences). Full length (FL) *yrt* refers to the beta isoform, which includes all exons of the *yrt* gene (Laprise et al., 2006). The resulting plasmids allowed for expression of the following 3xFLAG-tagged (N-terminus) Yrt proteins: ΔFERM (a.a. 1-56/346-972); VRΔPBM (a.a. 406-969), BH1KR>D (K124D, R132D, K134D); BH2K>D (K285D, K288D); BH2K>D,K>A (K285D, K288D, K292A, K293A). Genome Prolab (Sherbrooke, Canada) injected plasmids in *D. melanogaster* embryos carrying an attP docking site (Groth et al., 2004). Specifically, transgenes were inserted on the third chromosome (68E1; Bloomington *Drosophila* Stock Center (BDSC) line 24485, which was generated by the Basler’s group) or the second chromosome (25C6; line produced by Genome Prolab). The other constructs and fly lines used in this study were previously described (Gamblin et al., 2014; Gamblin et al., 2018). We employed the UAS-GAL4 system to express transgenes (Brand et al., 1994) using the da-GAL4 driver line (Wodarz et al., 1995). Embryo staging was achieved according to the Atlas of *Drosophila* Development (Hartenstein, 1993). Embryo sex was not determined, so both males and females were used indiscriminately. Plasmids and fly lines can be obtained by contacting the corresponding author.

### Cell culture and transfection

S2 cells, which were obtained from ATCC and used as is, were grown at room temperature in Schneider’s medium (Wisent Inc.) complemented with 10% fetal bovine serum (heat inactivated; Wisent Inc.), 50 units/ml penicillin, and 50 μg/ml streptomycin (Life Technologies). S2 cells were transfected at 60% confluency with FuGENE® (Promega) with selected pUASTattB-based constructs together with pAct5c-GAL4. When indicated (24 h post-transfection), S2 cells were treated with 10 μM ionomycin for 15 min, or 10 μM PAO and/or 1 μMμ wortmannin for 45 min.

### Immunofluorescence

In preparation for immunofluorescence on *D. melanogaster* embryos, the chorion was removed by a 5 min incubation in a solution of 3% sodium hypochlorite. Embryos were then washed with water, fixed for 8 min in 11% formaldehyde (diluted in 60 mM PIPES, 30 mM EGTA, 1.2 mM MgSO_4_) under a heptane phase (1:1), and rinsed with PBS. Embryos were devitellinized by vigorous agitation in methanol. Non-specific binding sites were blocked by a 1 h incubation in NGT (2% normal goat serum, 0.3% Triton X-100 in PBS). S2 cells were fixed 24 h post-transfection with 4% paraformaldehyde (in PBS) for 15 min, permeabilized with PBS-0,1% Triton X-100, and saturated for 1 h with a solution of 2% normal goat serum and 0.1% Triton X-100 in PBS.

The following primary antibodies were used overnight at 4°C: anti-FLAG (M2, Sigma), 1:250 ; anti-Yrt (Biehler et al., 2020), 1:500 ; anti-Lgl (d-300, Santa Cruz Technology), 1:100. Following incubation with primary antibodies, embryos and cells were washed three times for 20 min in PBS containing 0.3% or 0.1% Triton X-100 (PBT), respectively, and incubated for 1 h at room temperature with Cy3- or Alexa Fluor 488-conjugated secondary antibodies (1:400). Unbound secondary antibodies were washed-out with PBT (3 times 20 min). Embryos and cells were mounted in Vectashield mounting medium (Vector Labs) and imaged by confocal microscopy. Nuclei were stained with 4’,6-diamidino-2-phénylindole (DAPI).

### Western blotting

Dechorionated embryos were homogenized in lysis buffer (50 mM Tris-HCl pH 7.4, 150 mM NaCl, 1 mM EDTA, 1% Triton X-100, 0.1 mM sodium orthovanadate, 0.1 mM phenylmethylsulfonyl fluoride, 1 mM NaF, 10 μg/ml aprotinin, 10 μg/ml leupeptin and 0.7 µg/ml pepstatin) at 4°C and processed for SDS-PAGE and western blotting as described (Laprise et al., 2002). The following primary antibodies were used: anti-FLAG M2 (Sigma-Adrich) 1:4000; anti-GFP (Roche) 1:3000; anti-HA (BioLegend); anti-Actin (Novus Biologicals), 1:5000. Secondary antibodies were conjugated to HRP (GE Healthcare) and diluted 1:2000. TrueBlot ULTRA (Rockland) secondary antibodies were used for immunoprecipitation experiments (1:1000).

### Protein purification and lipid binding assays

GST and GST-FERM expression in BL21 (DE3) cells was induced by adding 0.1 mM IPTG to bacterial cultures (O.D. of 0.6 at 600 nm) for 16 h at 16°C. Proteins were then purified as previously described (Maity et al., 2013). PIP Strips membranes (P-6001; Echelon Biosciences) were saturated with 3% fatty acid free BSA (Sigma-Aldrich A8806) in PBS-Tween-20 0.1% for 1 h at room temperature. Membranes were then incubated with 1 µg of GST, GST-FERM, GST-VR, or PI(4,5)P2 Grip (GST-PLC-d1-PH; Echelon Biosciences) following Echelon Biosciences’ protocol. Protein-lipid interactions were revealed by western blot using an anti-GST antibody (Santa Cruz Biotechnology).

### Analysis of Yrt sequence

Wild type and mutated sequences of the Yrt FERM domain were analyzed with the BH-search algorithm [https://helixweb.nih.gov/bhsearch/ (Brzeska et al., 2010)].

### Quantification of membrane and cytoplasmic fluorescence intensity

Quantification of fluorescence signal intensity was performed with FIJI (Schindelin et al., 2012). In fly embryos, the polygon selection tool was used to generate a box, which was used to measure both membrane and cytoplasmic intensity in each cell and the mean membrane/cytoplasm ratio was determined. Ratios were determined in a total of 15 individual cells for each construct (5 non-adjacent cells/embryo; 3 different embryos randomly selected from different experiments). Sample size was empirically determined.

The freehand selection tool was used to individually select the whole membrane or cytoplasm in S2 cells. This method was chosen, as the signal intensity was often irregular along the membrane in S2 cells in contrast to what was observed *in vivo*. Total signal intensity (Raw Integrated Density; RID) and the area of each box were recorded, and the membrane/cytoplasm ratios were calculated as follow:

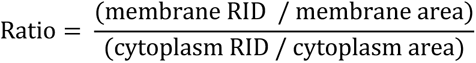

Fluorescence ratio was quantified in 39 to 62 individual cells for each construct (10 to 23 cells per experiment from 3-4 different experiments). The targeted sample size for S2 cells experiments was about 45 cells, as we devised it would show sample variability and be sufficient to meet the normal distribution assumption (true for samples treated with PAO and/or Wort).

### Viability test

One hundred newly laid embryos of each genotype were collected and placed on an apple plate at 25°C for 72 h (this experiment was performed in triplicate). Larvae and dead embryos were then scored, and the ratio of larvae on larvae plus dead embryos x 100 represents the hatching percentage. This sample size has been regularly used in our lab and allowed us to detect mild lethality phenotypes (Gamblin et al., 2018).

### Immunoprecipitations

HA immunoprecipitations were performed as previously described (Gamblin et al., 2018). For GFP-Trap, S2 cells were homogenized in lysis buffer at 4°C (50 mM Tris-HCl pH 7.4, 150 mM NaCl, 1 mM EDTA, 1% Triton X-100, 0.1 mM sodium orthovanadate, 0.1 mM phenylmethylsulfonyl fluoride, 1 mM NaF, 10 μg/ml aprotinin, 10 °C g/ml leupeptin and 0.7 µg/ml pepstatin. Cell lysates were centrifuged (17,000 g, 10 min at 4°C) to remove insoluble cell fragments. S2 cells lysates containing 2 mg of total proteins were precleared for 1.5 h under agitation at 4 °C with 15 µl of sepharose CL-4B beads (Sigma-Adrich). Supernatants were then incubated with 15 µl of GFP-Trap beads (ChromoTek) for 1.5 h at 4°C under agitation. Beads were recovered by centrifugation and washed 5 times with lysis buffer. Immunocomplexes were eluted with Laemmli’s buffer.

### Statistical analysis

Statistical analyses were performed with Prism 8 (GraphPad Software). Membrane-to-cytoplasmic ratio were presented as violin plots showing all individual measures as well as median and quartiles. As the data from the different groups were either not from a normal distribution and/or the sample size was relatively small, the differences between individual groups were analyzed using the non-parametric Kruskal-Wallis test and post-hoc Dunn’s multiple comparison test, all compared to FLAG-Yrt FL or DMSO (control conditions). Adjusted *p*-values <0.05 were considered statistically significant. In the case of ionomycin treatment in S2 cells, a non-parametric Mann-Whitney test, two-tailed with α=0.05 was performed to assess differences between treatments (as the data were not from a normal distribution). For viability test, hatching data were presented as means ± SEM. Differences between individual groups were analyzed by using Fisher’s exact test (95% confidence intervals, two-tailed). The significance threshold was adjusted in regards of the multiple comparisons with a Bonferroni correction (8 comparisons). *P*-values <0.00625 were considered statistically significant.

## Acknowledgments

We acknowledge Konrad Basler, Nicolas Bisson, Sébastien Carréno, Ulrich Tepass, and the Bloomington *Drosophila* Stock Center for reagents. Flybase was used as an important database for this work. This work was supported by operating grants from the Canadian Institute of Health Research (CIHR) to P.L. (PJT-185951) and M.L. (PJT-178198). F.C. is supported by a scholarship from FRQ-S, and C. B. is a postdoctoral Europe Horizon MSCA-PF grantee.

## Author contributions

P.L. conceived the project. C.G., C.A., F.C., F.P.P. and N.M. performed experiments and/or prepared reagents and contributed to data analysis. C.B. trained C.A. and contributed to initial *in vivo* analyses. A.J. performed statistical analysis and assembled figures with C.G. and C.A. P.L. wrote the manuscript, which was proofread by co-authors. P.L. and M.L. secured funding.

---

**Figure S1.**
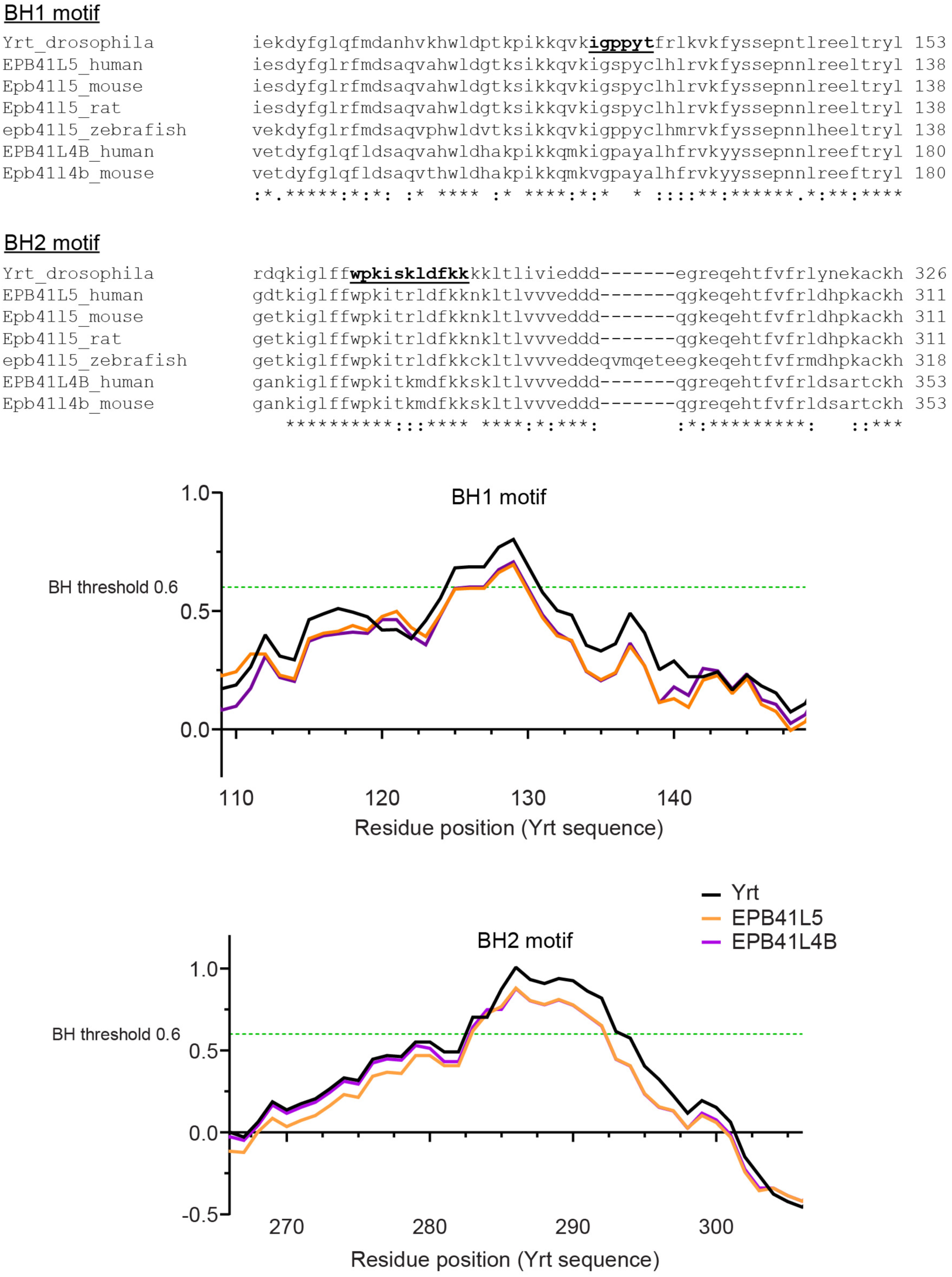
BH motifs in Yrt sequence are evolutionarily conserved. Alignment of the BH motifs of Yrt with the sequence of its two orthologs, namely EPB41LS and EPB41L4B. Bottom panels show the BH score for Yrt sequence (black curve), or human EPB41L4B (purple curve) and EPB41LS (orange curve) sequences. The BH1/2 scores of EPB41L4B and EPB41LS go beyond the threshold of 0.6 even though their respective sequence diverges from Yrt sequence to some extent. Overall, these data suggest that Yrt orthologs also interact with the membrane using, among others, the conserved BH1 and BH2 motifs.

## REFERENCES

Bailey, M. J. and Prehoda, K. E. (2015). Establishment of Par-Polarized Cortical Domains via Phosphoregulated Membrane Motifs. Dev Cell 35, 199–210.

Baines, A. J. (2006). A FERM-adjacent (FA) region defines a subset of the 4.1 superfamily and is a potential regulator of FERM domain function. BMC Genomics 7, 85.

Banerjee, S., Sousa, A. D. and Bhat, M. A. (2006). Organization and function of septate junctions: an evolutionary perspective. Cell Biochem Biophys 46, 65–77.

Barnett, K. C., Coronas-Serna, J. M., Zhou, W., Ernandes, M. J., Cao, A., Kranzusch, P. J. and Kagan, J. C. (2019). Phosphoinositide Interactions Position cGAS at the Plasma Membrane to Ensure Efficient Distinction between Self- and Viral DNA. Cell 176, 1432–1446 e11.

Bieber, A. J., Snow, P. M., Hortsch, M., Patel, N. H., Jacobs, J. R., Traquina, Z. R., Schilling, J. and Goodman, C. S. (1989). Drosophila neuroglian: a member of the immunoglobulin superfamily with extensive homology to the vertebrate neural adhesion molecule L1. Cell 59, 447–60.

Biehler, C., Rothenberg, K. E., Jette, A., Gaude, H. M., Fernandez-Gonzalez, R. and Laprise, P. (2021). Pak1 and PP2A antagonize aPKC function to support cortical tension induced by the Crumbs-Yurt complex. Elife 10.

Biehler, C., Wang, L. T., Sevigny, M., Jette, A., Gamblin, C. L., Catterall, R., Houssin, E., McCaffrey, L. and Laprise, P. (2020). Girdin is a component of the lateral polarity protein network restricting cell dissemination. PLoS Genet 16, e1008674.

Brand, A. H., Manoukian, A. S. and Perrimon, N. (1994). Ectopic expression in Drosophila. Methods Cell Biol 44, 635–54.

Brzeska, H., Guag, J., Remmert, K., Chacko, S. and Korn, E. D. (2010). An experimentally based computer search identifies unstructured membrane-binding sites in proteins: application to class I myosins, PAKS, and CARMIL. J Biol Chem 285, 5738–47.

Buckley, C. E. and St Johnston, D. (2022). Apical-basal polarity and the control of epithelial form and function. Nat Rev Mol Cell Biol.

Dong, W., Zhang, X., Liu, W., Chen, Y. J., Huang, J., Austin, E., Celotto, A. M., Jiang, W. Z., Palladino, M. J., Jiang, Y. et al. (2015). A conserved polybasic domain mediates plasma membrane targeting of Lgl and its regulation by hypoxia. J Cell Biol 211, 273–86.

Ehret, T., Heissenberg, T., de Buhr, S., Aponte-Santamaria, C., Steinem, C. and Grater, F. (2023). FERM domains recruit ample PI(4,5)P(2)s to form extensive protein-membrane attachments. Biophys J 122, 1325–1333.

Gamblin, C. L., Hardy, E. J., Chartier, F. J., Bisson, N. and Laprise, P. (2014). A bidirectional antagonism between aPKC and Yurt regulates epithelial cell polarity. J Cell Biol 204, 487–95.

Gamblin, C. L., Parent-Prevost, F., Jacquet, K., Biehler, C., Jette, A. and Laprise, P. (2018). Oligomerization of the FERM-FA protein Yurt controls epithelial cell polarity. J Cell Biol.

Gassama-Diagne, A., Yu, W., ter Beest, M., Martin-Belmonte, F., Kierbel, A., Engel, J. and Mostov, K. (2006). Phosphatidylinositol-3,4,5-trisphosphate regulates the formation of the basolateral plasma membrane in epithelial cells. Nat Cell Biol 8, 963–70.

Groth, A. C., Fish, M., Nusse, R. and Calos, M. P. (2004). Construction of transgenic Drosophila by using the site-specific integrase from phage phiC31. Genetics 166, 1775–82.

Hartenstein, V. (1993). Atlas of Drosophila development. Plainview, N.Y.: Cold Spring Harbor Laboratory Press.

Hoover, K. B. and Bryant, P. J. (2002). Drosophila Yurt is a new protein-4.1-like protein required for epithelial morphogenesis. Dev Genes Evol 212, 230–8.

Krahn, M. P. (2020). Phospholipids of the Plasma Membrane - Regulators or Consequence of Cell Polarity? Front Cell Dev Biol 8, 277.

Laprise, P., Beronja, S., Silva-Gagliardi, N. F., Pellikka, M., Jensen, A. M., McGlade, C. J. and Tepass, U. (2006). The FERM protein Yurt is a negative regulatory component of the Crumbs complex that controls epithelial polarity and apical membrane size. Dev Cell 11, 363–74.

Laprise, P., Chailler, P., Houde, M., Beaulieu, J. F., Boucher, M. J. and Rivard, N. (2002). Phosphatidylinositol 3-kinase controls human intestinal epithelial cell differentiation by promoting adherens junction assembly and p38 MAPK activation. J Biol Chem 277, 8226–34.

Laprise, P., Lau, K. M., Harris, K. P., Silva-Gagliardi, N. F., Paul, S. M., Beronja, S., Beitel, G. J., McGlade, C. J. and Tepass, U. (2009). Yurt, Coracle, Neurexin IV and the Na(+),K(+)-ATPase form a novel group of epithelial polarity proteins. Nature 459, 1141–5.

Laprise, P., Paul, S. M., Boulanger, J., Robbins, R. M., Beitel, G. J. and Tepass, U. (2010). Epithelial polarity proteins regulate Drosophila tracheal tube size in parallel to the luminal matrix pathway. Curr Biol 20, 55–61.

Maity, R., Pauty, J., Krietsch, J., Buisson, R., Genois, M. M. and Masson, J. Y. (2013). GST-His purification: a two-step affinity purification protocol yielding full-length purified proteins. *J Vis Exp*, e50320.

Martin-Belmonte, F., Gassama, A., Datta, A., Yu, W., Rescher, U., Gerke, V. and Mostov, K. (2007). PTEN-mediated apical segregation of phosphoinositides controls epithelial morphogenesis through Cdc42. Cell 128, 383–97.

Ohata, S., Aoki, R., Kinoshita, S., Yamaguchi, M., Tsuruoka-Kinoshita, S., Tanaka, H., Wada, H., Watabe, S., Tsuboi, T., Masai, I. et al. (2011). Dual roles of Notch in regulation of apically restricted mitosis and apicobasal polarity of neuroepithelial cells. Neuron 69, 215–30.

Sparks, L. M., Moro, C., Ukropcova, B., Bajpeyi, S., Civitarese, A. E., Hulver, M. W., Thoresen, G. H., Rustan, A. C. and Smith, S. R. (2011). Remodeling lipid metabolism and improving insulin responsiveness in human primary myotubes. PLoS One 6, e21068.

Tepass, U. (2009). FERM proteins in animal morphogenesis. Curr Opin Genet Dev 19, 357–67.

Wodarz, A., Hinz, U., Engelbert, M. and Knust, E. (1995). Expression of crumbs confers apical character on plasma membrane domains of ectodermal epithelia of Drosophila. Cell 82, 67–76.

Wymann, M. P., Zvelebil, M. and Laffargue, M. (2003). Phosphoinositide 3-kinase signalling--which way to target? Trends Pharmacol Sci 24, 366–76.

